# Inferring a cell’s capabilities from omics data with ImmCellFie

**DOI:** 10.1101/2022.11.16.516672

**Authors:** Helen O. Masson, David Borland, Jason Reilly, Adrian Telleria, Shalki Shrivastava, Matt Watson, Luthfi Bustillo, Zerong Li, Laura Capps, Benjamin P. Kellman, Zachary A. King, Anne Richelle, Nathan E. Lewis, Kimberly Robasky

## Abstract

ImmCellFie is a user-friendly, web-based platform for comprehensive analysis of metabolic functions inferred from transcriptomic or proteomic data. It enables researchers to leverage the powerful mechanistic insight provided by complex genome-scale metabolic models with little to no bioinformatics training required. The platform has been integrated with a series of useful tools and richly annotated scientific visualizations for interactive exploration by the user. ImmCellFie pushes beyond simple statistical enrichment and incorporates complex biological mechanisms to quantify cell activity.

Graphical abstract

## Before you begin

### Overview

Life functions through complex interdependencies between genes, proteins, and metabolites. The advent of high-throughput omic technologies have enabled researchers to comprehensively monitor these individual molecule-types; however, interpretation of such omics experiments remains challenging due to the aforementioned interdependence between molecular components. While statistical methods are invaluable for identifying sets of genes involved in a given phenotype, it remains difficult to obtain and quantify mechanisms underlying a cell’s functions from only enriched pathway and ontology terms.

Genome-scale metabolic networks (Feist *et al*., 2009; Lewis, Nagarajan and Palsson, 2012) provide systems biologists with tools to analyze genome-scale omics datasets. These reconstructed networks directly link genotype to phenotype via a mathematical knowledge base of all metabolic reactions within an organism. Unfortunately, working with these models requires specialized training and extensive effort to deploy. There is a need for more accessible tools that allow any researcher to obtain similar mechanistic insights from omics data. To this end we developed CellFie (Richelle *et al*., 2021), an approach that leverages 195 metabolic tasks identified in mammalian cells using genome-scale metabolic models, and enables one to overlay transcriptomic and proteomic data to quantify activity changes in each metabolic task. These metabolic tasks are organized into a hierarchical structure of systems and subsystems, grouping tasks based on their biological functions. Using the CellFie platform, researchers have studied the metabolic impacts of many systems, including perturbing renal drug transporters (Granados *et al*., 2021), and identifying metabolic tasks perturbed in Alzheimer’s disease (Richelle *et al*., 2021).

ImmCellFie (immcellfie.renci.org), presented here, was developed as a web-based platform to house the CellFie package, enabling any scientist, regardless of computational background, to directly predict, quantify, and visualize how changes in omics experiments correspond to the metabolic functions of cells or tissues. Users can easily upload proteomic or transcriptomic data and use the provided suite of analytical and interactive visualization tools to explore the metabolic activity of their dataset. The protocol presented here describes how to use ImmCellFie to infer metabolic capabilities from omics data and visualize the complex mechanisms driving metabolic phenotypes. RNA-Seq data from the Human Protein Atlas (HPA) is used as a case-study for this goal.

ImmCellfie uses genome-scale metabolic models (GeMs) to quantify metabolic task activity from omics data. The algorithm precomputes metabolic pathway activities involving thousands of metabolic reactions (Sigurdsson *et al*., 2010; Hefzi *et al*., 2016; Swainston *et al*., 2016) to find gene modules (“metabolic tasks”) that collectively consume a metabolite to produce a final metabolic product of interest (Richelle *et al*., 2019). When transcriptomic or proteomic data are overlaid on the precomputed modules, pathway activities are estimated for each sample. Complementing current enrichment analyses, CellFie quantifies a gene set’s impact on the cell by mathematically defining the function of all genes and determining the expression requirements to accomplish metabolic tasks (see the *Understanding metabolic task calculation* for detailed description of task score calculations). ImmCellfie supports several model organisms including human, mouse, rat, and Chinese hamster (CHO). In this protocol we show how users can conduct rich analyses of omics data with the click of a button, by uploading data, submitting customized jobs, and exploring the results with useful tools and richly annotated scientific visualizations including heatmaps, Voronoi treemaps, and interactive pathway maps.

### Software download and prerequisites

ImmCellfie can either be operated via the web dashboard (immcellfie.renci.org) or through an OpenAPI application programming interface (API). The API can be used for running data in batch and for integrating with notebooks (e.g., Jupyter, Google Colab). For documentation on interfacing with the API, see http://immcellfie.renci.org:8081/docs. The protocol presented here describes the steps for running ImmCellFie via the web dashboard. Here, we provide specific instructions for preprocessing the HPA example expression data in R prior to running ImmCellFie. However, this preprocessing step can be performed using any programming language and/or online gene-mapping web tool as long as the end result meets ImmCellFie formatting requirements. The easiest option to follow the preprocessing steps outlined in this tutorial is to use RStudio. Both R and RStudio can be downloaded from https://www.r-project.org/ and https://rstudio.com/products/rstudio/download/, respectively.

### Understanding metabolic task calculation

Metabolic tasks are defined by the capacity to produce a defined list of output products when only a defined list of input substrates is available. For each available reference GeM, we define a list of tasks supported by the model. Then the reactions required to complete each task are identified. Based on the model’s gene-protein-reaction (GPR) rules, we link the task with genes contributing to the metabolic function. CellFie predicts the relative activity of each task using the expression level of the associated gene set by first attributing a gene score to each expressed gene in the model.

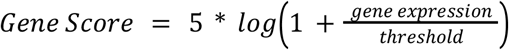

The computation of the Gene Score required one to define a threshold that specifies whether a gene is expressed or not. One can use multiple options to define the gene threshold (see section *Threshold Definition* for more details)(Richelle, Joshi and Lewis, 2019). Gene Scores are mapped to the GeM via parsing of the GPR rule associated with each reaction: select the minimum expression amongst all the genes associated with an enzyme complex (AND rule) and the maximum expression amongst all the genes associated with an isozyme (OR rule)(Jensen, Lutz and Papin, 2011). As a result, each reaction is assigned a reaction activity level (RAL) equal to the gene score of the main determinant of the reaction.

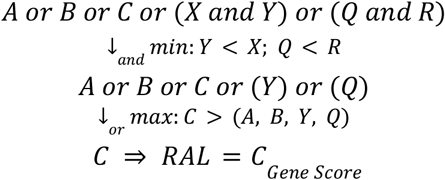

We account for the competition of promiscuous enzymes/genes (i.e. genes used in multiple reactions) by scaling each gene with a specificity score (S).

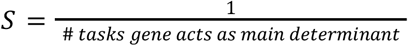

Finally, the metabolic task score (MTS) can be computed as the mean scaled reaction activity level.

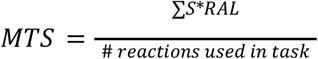

Some important housekeeping genes are always present at very low expression values, therefore metabolic tasks that will solely rely on these types of genes will always result in a low MTS. In contrast, some tasks are associated with highly expressed genes. This is the reason why tasks should not be compared within the same sample using the quantitative score value. To facilitate intra-sample comparison of tasks, we also modify this scoring approach to a binary form defining tasks as either active or inactive. To this end, a metabolic task is considered active if the average of its associated RAL is greater than 5*log(2).

### Defining gene thresholds

We propose two thresholding approaches to define which genes from the input expression data are considered “active”(Richelle, Joshi and Lewis, 2019).

1. Global approach, the threshold value is the same for all the genes. The global approach is mainly applied when only one sample is available (i.e. sample could be associated to a condition, a cell-type or a tissue) and/or no information is available in the literature to define expression threshold for a single gene. The user can define the global threshold using a value or a percentile
  a. value: the user provides a set value above which a gene will be considered active
  b. percentile: the threshold value will be defined based on a percentile (defined by the user) of the distribution of expression values for all the genes and across all samples of the user dataset
2. Local approach, the threshold value is gene-specific. The local approach is often applied when multiple samples are available as it enables a relative assessment of the activity of a gene across samples. Multiple options are available to compute the local thresholds:

a. mean: the threshold for a gene is defined as the mean expression value of this gene across all the samples, tissues, or conditions.
b. min-max mean: the threshold for a gene is determined by the mean of expression values observed for that gene among all the samples, tissues, or conditions, but the threshold :(i) must be higher or equal to a lower bound and (ii) must be lower or equal to an upper bound. The lower and upper bound can be defined using a value introduced by the user or based on a percentile of the distribution of expression values for all the genes and across all samples in the dataset. The MinMaxMean option ensures that genes with very low expression values across all the samples will never be considered as active and genes with very high expression across samples are always considered as active.

## Key resources table

| REAGENT or RESOURCE | SOURCE | IDENTIFIER |
| --- | --- | --- |
| Deposited data |  |  |
| RNA HPA blood cell sample gene data (rna_blood_cell_sample.tsv.zip) | HPA | https://www.proteinatlas.org/about/download |
| demo_expression.csv | This manuscript | n/a |
| demo_pheno.csv | This manuscript | n/a |

| Software and algorithms |  |  |
| --- | --- | --- |
| ImmCellFie | http://immcellfie.renci.org | This manuscript |
| R | R Core Team (2020). R: A language and environment for statistical computing | https://www.R-project.org/ |
| data.table | Matt Dowle and Arun Srinivasan (2021). data.table: Extension of `data.frame` | https://CRAN.R-project.org/package=data.table |
| dplyr | Hadley Wickham, Romain François, Lionel Henry and Kirill Müller (2021). dplyr: A Grammar of Data Manipulation | https://CRAN.R-project.org/package=dplyr |
| tidyr | Hadley Wickham and Maximilian Girlich (2022). tidyr: Tidy Messy Data | https://CRAN.R-project.org/package=tidyr |
| AnnotationDbi | Hervé Pagès, Marc Carlson, Seth Falcon and Nianhua Li (2020). AnnotationDbi: Manipulation of SQLite-based annotations in Bioconductor | http://doi.org/10.18129/B9.bioc.AnnotationDbi |
| org.Hs.eg.db | Marc Carlson (2020). org.Hs.eg.db: Genome wide annotation for Human | http://doi.org/10.18129/B9.bioc.org.Hs.eg.db |

## Step-by-step method details

### Create an account with ImmCellFie

**Timing:** 30 seconds

User accounts store imported datasets and CellFie results that users can return to and share with colleagues.

1. Navigate to the ImmCellFie home page: http://immcellfie.renci.org
2. Click on the ‘User’ tab at the top of the page.
3. Enter your email address as username.

#### CRITICAL

ImmCellFie accounts are NOT password protected. Uploaded data will be publicly available to other ImmCellFie users with access to your username. Please ensure that no Protected Health Information (PHI) is uploaded. Also take care with uploading unpublished data. For protected data we recommend locally installing and running CellFie.

### Preparing ImmCellFie input files

**Timing:** 5-10 minutes

ImmCellFie requires a specific format for input data. Expression data must meet the following format requirements: i) entrez gene IDs in the first column, ii) wide format (each subsequent column contains the expression data for individual samples), iii) uploaded as csv, and iv) optional header in the first row with sample names. Users also have the option to include a properties file containing phenotypic information to group samples. This file must: i) include a header with property names (e.g. sex, cell type, treatment, etc.), ii) match row order to the sample order (columns) in the expression data, iii) uploaded as csv. Input data needs to be pre-processed to meet these formatting requirements. The following steps outline how the HPA blood expression data was pre-processed in R.

4. Download the expression data from HPA as a zip folder and save into a directory of your choice.
5. Open RStudio.
6. Load the required library packages.

~~~
  > library(data.table)
  > library(dplyr)
  > library(tidyr)
  > library(AnnotationDbi)
  > library(org.Hs.eg.db)
~~~
7. Load HPA data.

~~~
  > blood.data <- fread(paste(‘unzip -p’,
  ‘Blood_Atlas_Data/rna_blood_cell_sample.tsv.zip’))
~~~ **Note**: We have created a folder called ‘Blood_Atlas_Data’ as our directory of choice in this example. Ensure that your RStudio working directory is set to the same location that houses this Blood_Atlas_Data folder.
8. Generate expression matrix compatible with CellFie.

a. Isolate expression data and reshape the data such that rows are samples and columns are genes

~~~
> TPM.matrix <- blood.data[, c(“ENSG ID”, “Sample ID”, “TPM”)] %>%
       spread(‘Sample ID’, TPM)
~~~
b. Convert Ensembl gene IDs to entrez IDs.

~~~
> gene.map <- AnnotationDbi::select(org.Hs.eg.db,
      keys=TPM.matrix$‘ENSG ID’,
      columns=c(“ENTREZID”),
      keytype=“ENSEMBL”) %>% na.omit()
> TPM.matrix.entrez <- left_join(gene.map, TPM.matrix,
      by=c(“ENSEMBL”=“ENSG ID”)) %>% dplyr::select(-ENSEMBL)
~~~
c. Aggregate duplicate genes by max value.

~~~
> TPM.entrez.max <- aggregate(TPM.matrix.entrez,
       by=list((TPM.matrix.entrez[,1])),max) %>%
       dplyr::select(-Group.1)
~~~
d. Add pseudo count.

~~~
> TPM.entrez.pseudo <- cbind(ENTREZID=TPM.entrez.max$ENTREZID,
       TPM.entrez.max[, 2:ncol(TPM.entrez.max)] + 0.00001)
~~~
9. Generate properties data matrix (phenotypic information for the samples).

~~~
 > pheno.data <- dplyr::select(blood.data, c(“Sample ID”, “Donor”,
        “Immune cell”)) %>% distinct()
~~~
10. Export files.

~~~
  > fwrite(TPM.entrez.pseudo, “demo_expression.csv”)
  > fwrite(pheno.data, “demo_pheno.csv”)
~~~

#### Note

We have provided an R Markdown (RMD) file with the complete code to preprocess these files (Supplementary File 1). The end result of these steps is the generation of two CSV files: demo_expression.csv and demo_pheno.csv. Users are free to use any platform to process files to obtain CSV files. If users wish to avoid using R in this tutorial, they can skip this step and use the provided processed files (Supplementary Files 2-3) to follow along with the rest of the tutorial.

#### Alternatives

CSV files can also be generated using spreadsheet programs, such as Microsoft Excel, to make tables that match the layouts of demo_expression.csv and demo_pheno.csv, and save the spreadsheets in CSV format.

### Upload data to ImmCellFie

**Timing:** seconds

ImmCellFie allows users to upload data in two ways: i) locally from a personal computer and ii) import data directly from ImmuneSpace.org. Here we will be uploading the expression data and properties document saved locally from the previous step. **<u>Troubleshooting 1</u>**

11. Click ‘Select data’ below your newly entered user name, or click on the ‘Data’ tab on the ImmCellFie Dashboard. Click on ‘Upload local files’. A pop-up will appear with detailed formatting instructions.
12. Upload the csv files generated from the previous steps: demo_expression.csv and demo_pheno.csv.
13. Annotate the data set as “ImmCellFie Demo” in the ‘Description (optional)’ box.
14. Click ‘Upload’ to submit the data set. A pop up will appear indicating that the data has been successfully uploaded to your account.

#### Alternatives

Users have the option to directly import data from the ImmuneSpace database: https://www.immunespace.org/. While a complete tutorial of the ImmuneSpace database is beyond the scope of this protocol, navigating their website is quite self-explanatory. To import data groups from your ImmuneSpace account, simply click on ‘ImmuneSpace’ rather than ‘Upload local files’ from the ImmCellFie Data page and follow the detailed instructions (hover over the question marks) to connect your custom ImmuneSpace group with ImmCellFie. For this you will need to obtain an API Key from ImmuneSpace and identify Group Labels for the data you wish to use.

### Run CellFie algorithm

**Timing:** 9 minutes (can be longer for larger datasets) **<u>Troubleshooting 2</u>**

The CellFie algorithm presents a powerful approach to predict the activity of many metabolic functions from transcriptomic and proteomic data. This framework facilitates phenotypic interpretation from these complex omic data types by comprehensively quantifying the propensity of a cell line or tissue to express the molecular components underlying a set of metabolic functions. Metabolic task scores can be computed based on any type of transcriptomic (e.g., microarray or RNA-Seq, bulk or single cell) or proteomic dataset for CHO, human, rat, or mouse. Additionally, users can select various types of gene-expression thresholding methods.

15. The previous step merely uploaded data to your ImmCellFie account. Next you will load the dataset as input to the CellFie algorithm (Figure 2A).

a. If the pop-up window is still open, you can click on the ‘Load’ button at the bottom right. Alternatively, you can navigate to the Data page from the ImmCellFie Dashboard and click the ‘Load’ button to the right of the ImmCellFie Demo dataset. **Note**: The dashboard should now read ‘ImmCellFie Demo’ under ‘Input’.
16. Click ‘Analyze’ from the ImmCellFie Dashboard, and select ‘CellFie’ from the drop-down menu. Alternatively, users may click on ‘Analyze’, followed by ‘CellFie’. (Figure 2B)
17. The left side of the page contains the necessary parameters for running a CellFie analysis. Select ‘human’ from the ‘Organism’ drop-down menu.

a. This will automatically select the correct human model ‘recon v2.2’.
18. CellFie provides two thresholding strategies to determine the set of “active” genes in the input dataset. The global approach is mainly applied when only one sample is available, therefore we will be using the gene-specific local gene thresholding approach in this example. Select ‘local’ from the ‘Threshold type’ drop-down menu and keep the default local thresholding parameters. See *Defining gene thresholds* section for detailed descriptions of the thresholding approaches.

a. Percentile or value: percentile
b. Local threshold type: min-max mean
c. Low percentile: 25
d. High percentile: 75
19. Add the optional description “demo-results” in the ‘Description’ box. **<u>Troubleshooting 2</u>**
20. Click ‘Run CellFie’. A pop-up window will appear with an estimate of how long the run will take. After exiting the pop-up, the ImmCellFie dashboard will continue to display a tiny spinning wheel on the far right to indicate that the run is computing. (Figure 2C)
21. When the run is complete, click ‘Load’ from the popup window to load the CellFie results. Alternatively, users can load results by navigating to the Data page from the ImmCellFie Dashboard and use the load button to the right of the CellFie run. **<u>Troubleshooting 3</u>** **Note:** The dashboard should now read ‘demo-results’ under ‘Result’.

**Figure 1:**
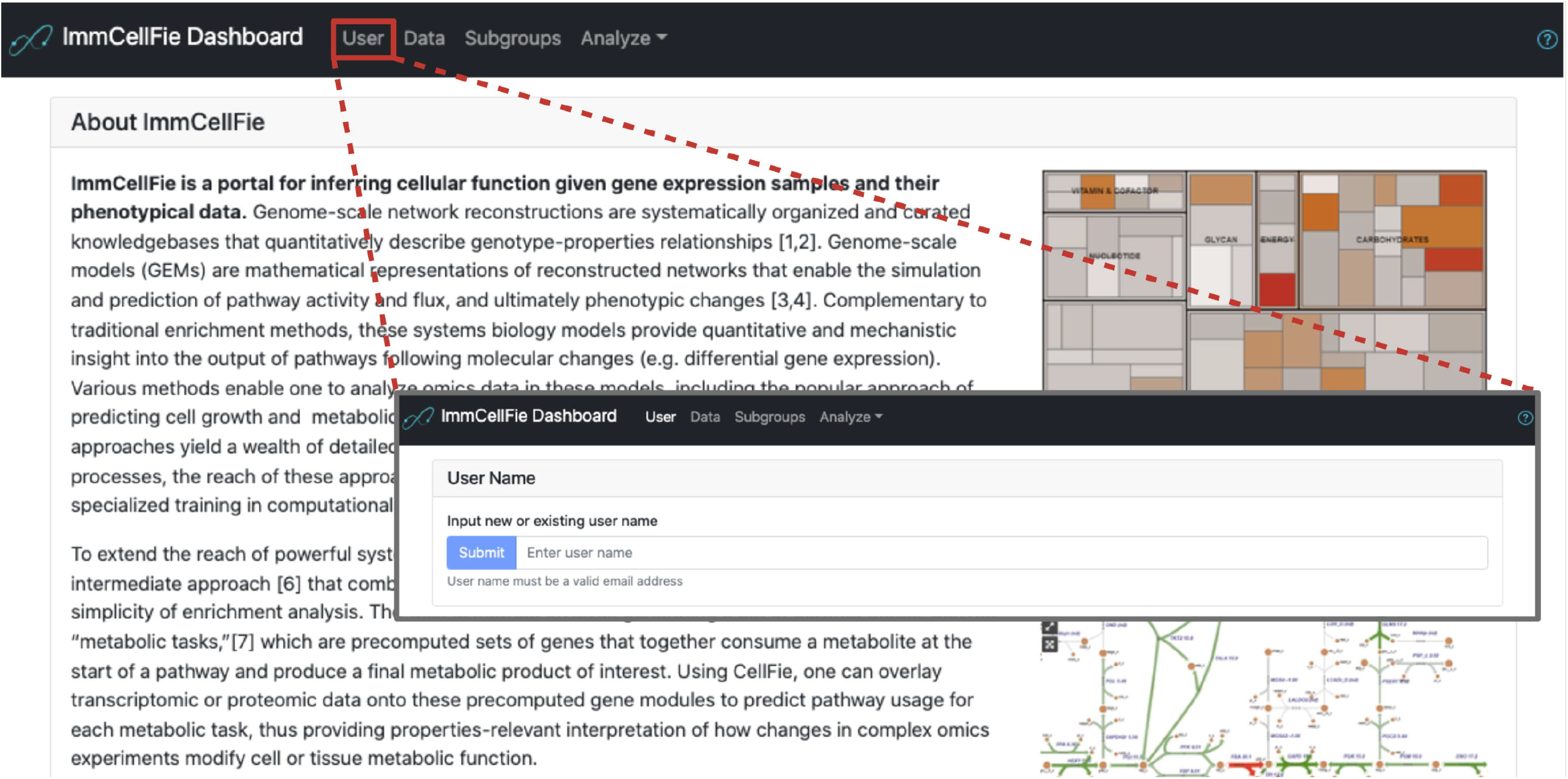
Setting up a user account with ImmCellFie. Screenshots showing how users create an account (or login in with a pre-existing account) with ImmCellFie.

**Figure 2:**
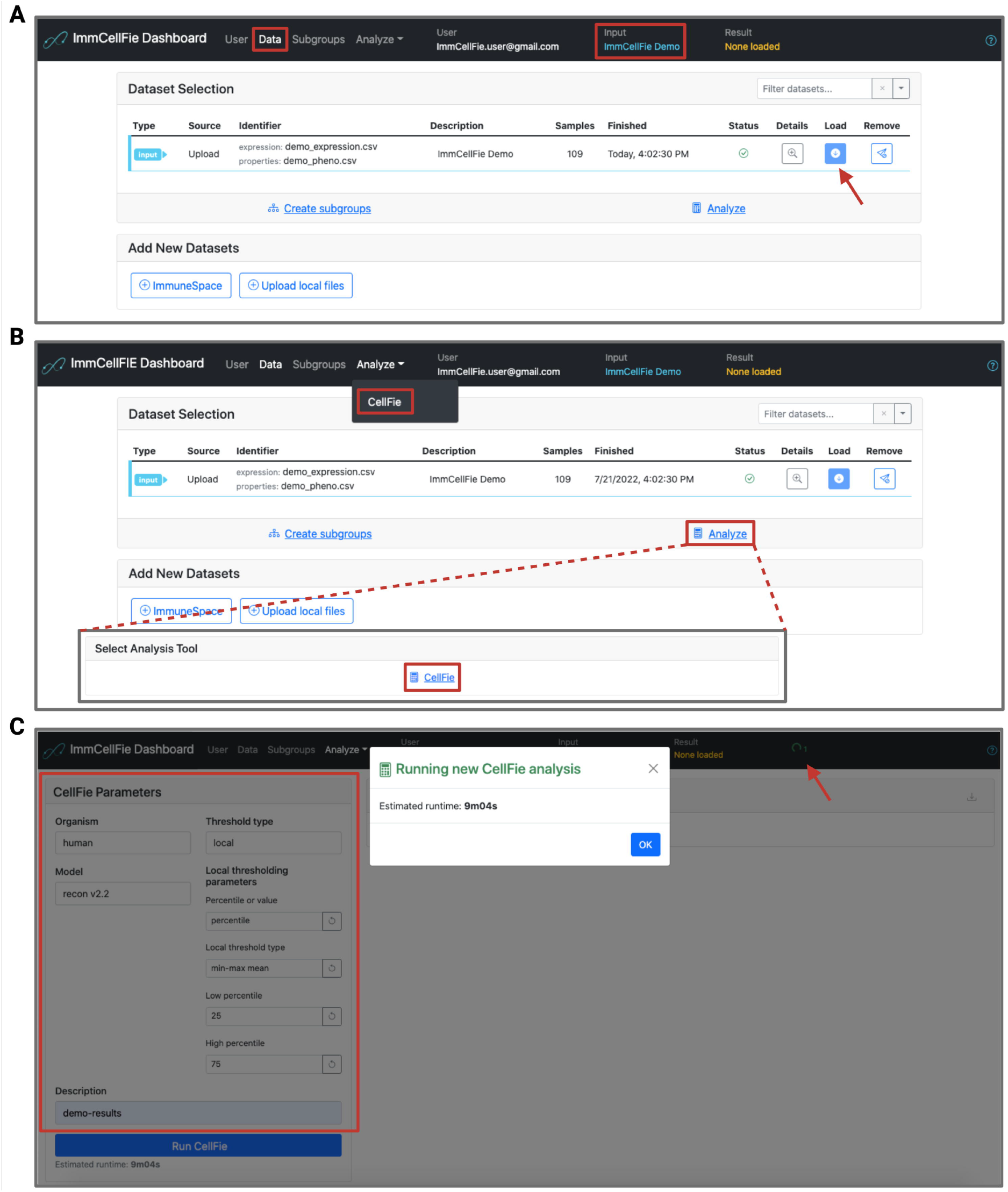
Setting up a CellFie run. Series of screenshots demonstrating various steps of setting up a CellFie run in ImmCellFie. A) Load input data; B) Two methods of navigating to the CellFie setup page; C) Setting CellFie parameters (red box) and starting a run.

### Download CellFie results

**Timing:** seconds

Users can download the raw output of a CellFie run. These results can be further analyzed by the user using their platform of choice. The raw output of CellFie includes 4 files (Table 1). Briefly, the main output of CellFie is a matrix of metabolic task scores (MTSs). The MTS output of CellFie provides a quantification of each metabolic function within the samples. Due to the nature of how MTSs are calculated, scores should not be compared across tasks (within a sample) but only across samples. To facilitate intra-sample comparison of tasks, CellFie also calculates MTSs in a binary form, which simply applies a task-specific threshold to determine whether the task is active or not.

22. Use the ‘Downloads’ button 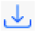 at the top of the CellFie Results track to batch-download all the results.

**Table 1.** Description of CellFie results included in batch-download.

| Results File | Description |
| --- | --- |
| score.csv | Matrix of task scores. Rows correspond to individual tasks and columns correspond to samples. |
| scoreBinary.csv | Matrix of binary task score activity (1=active, 0 = inactive). Rows correspond to individual tasks and columns correspond to samples. |
| taskInfo.csv | Table linking tasks to their respective metabolic pathways systems and subsystems. This is used to map rows of the score matrices to their respective tasks. |
| detailScoring.csv | Detailed description of genes and reactions used to calculate task scores for every sample:<br>• Column 1: ID of the sample<br>• Column 2: ID of the metabolic task<br>• Column 3: Task score for this sample<br>• Column 4: Binary task score for this sample<br>• Column 5: Essential reaction associated with this task<br>• Column 6: Expression score associated with the reaction listed in column 5 (i.e. RAL -Reaction Activity Level)<br>• Column 7: Gene used to determine the expression of the reaction in column 5<br>• Column 8: Original expression value of the gene listed in column 7 Information is concatenated horizontally therefore columns 9-16 will contain information for sample 2, columns 17-24 will contain information for sample 3, etc. |

#### Note

The score matrices do not contain row or column headers. Row labels for the score matrices (score.csv and scoreBinary.csv) are provided in a separate matrix file (taskInfo.csv), and the column order of the score matrices correspond to the order of samples provided in the input expression file. Appending row labels and headers can be easily done in any spreadsheet software (e.g., Microsoft Excel, Google Sheets, etc.).

### Use properties data to generate subgroups based on cell type

**Timing:** seconds to minutes

Users can generate subgroups based on information (columns) provided in the properties document. This enables users to explore meaningful comparisons between various groups of samples.

23. Click ‘Create subgroups’ from the ‘Select Subgroups to Compare’ box on the bottom left of the screen. Alternatively, users can navigate to the Subgroups page from the ImmCellFie Dashboard.

a. This page contains statistics for each data grouping based on the columns provided in the properties document: Sample ID, Donor, and Immune cell. So far the only existing group is ‘All samples’.
24. Split the data into subgroups based on cell type provided in the properties document. (Figure 3A)

a. Click on the button below ‘Immune cell’ to split the data into subgroups. You should now see grouping statistics for each cell type in the dataset (MAIT T-cell, NK-cell, T-reg, basophil, etc.).
25. Group cell types into granulocytes, monocytes, dendritic cells, B-cells, and T-cell custom subgroups.

a. Click on ‘Add subgroup’ at the bottom of the Subgroups page
b. Name the subgroup “Granulocytes”
c. Use your mouse to select basophil, eosinophil, and neutrophil from the Immune Cell bar plot. (Figure 3B)
d. Repeat step a-c to create custom groups for i) “Monocytes”: classical monocyte, intermediate monocyte, and non-classical monocyte, ii) “Dendritic Cells”: myeloid DC and plasmacytoid DC, iii)”B cells”: memory B-cell and naive B-cell, and iv) “T Cells”: MAIT T-cell, T-reg, gdT-cell, memory CD4 T-cell, memory CD8 T-cell, naive CD4 T-cell, and naive CD8 T-cell.
26. Return the results page by clicking on ‘Analyze’ >> ‘CellFie’ from the ImmCellFie Dashboard.

**Figure 3:**
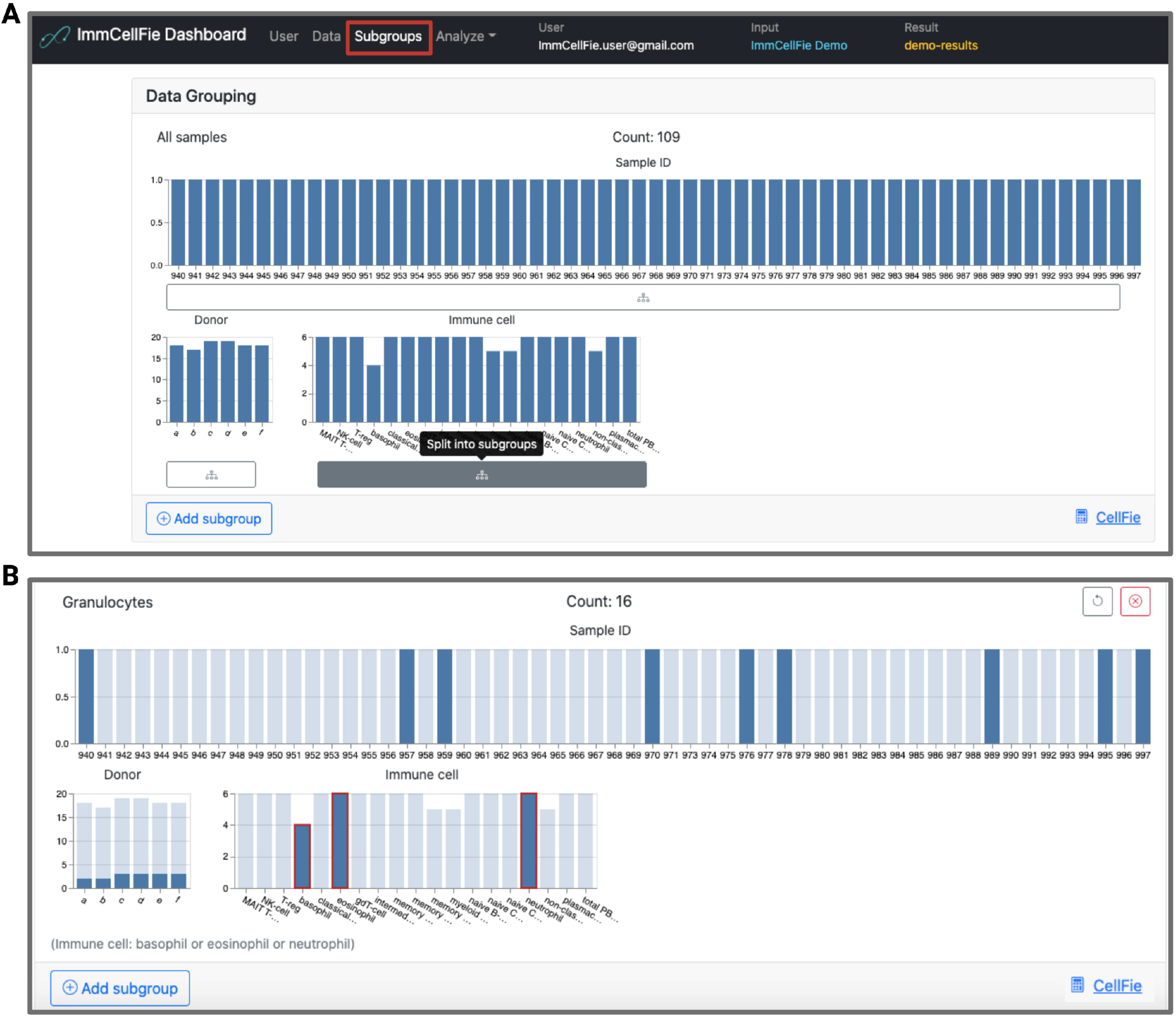
Creating subgroups based on immune cell type. Series of screenshots demonstrating how to create subgroups. A) Creating subgroups based on columns from the properties document upload; B) Example of creating a custom subgroup.

#### Note

Dividing the data into subgroups may take a few seconds. Do not click multiple times as this will generate multiple copies of the same groupings.

#### Note

Users can navigate to the Subgroups page from the ImmCellFie dashboard at any time to view and generate subgroups for comparison.

### Explore activity scores within the CellFie Hierarchy

**Timing:** minutes

CellFie tasks are organized into a hierarchical structure of system and subsystems, grouping tasks based on their biological functions. ImmCellFie provides various interactive tools for users to explore and visualize scores at various depths within the CellFie hierarchy.

27. Click the ‘Tree’ tab from the CellFie Results. (Figure 4)

a. By default the first two subgroups will be selected for comparison: MAIT T-cell and NK-cell.
b. This interactive tool shows the architecture of the Cellfie hierarchy linking tasks to their respective systems and subsystems, while including annotations of the score (continuous) and activity (binary) results. Users can click on various hierarchy elements to expand/shrink layers.
28. Users can adjust several parameters on the treemap:

a. Depth: refers to the system (Depth 1), subsystem (Depth 2), and task (Depth 3) levels within the CellFie hierarchy.
b. Subgroup: provides a drop-down menu allowing the user to visualize the mean score/activity of individual groups (MAIT T-cell OR NK-cell), or the difference between two groups (MAIT T-cell vs. NK-cell).
c. Color map: gives the user a choice of color schemes.
d. Search: Provides a drop-down menu where users can find and select specific elements within the CellFie hierarchy (e.g. “Krebs cycle - NADH generation”).
29. While the Tree tool provides a means for users to explore the CellFie hierarchy to clearly understand the architectural links between tasks-systems-subsystems, it does not generate publication-ready visualizations. The ImmCellFie Hierarchy tool however provides hierarchical visualizations in the form of a treemap, enclosure diagram, or Voronoi map. Click on the ‘Hierarchy’ tab in the CellFie Results.
30. Use the drop-down menus under ‘Select Subgroups to Compare’ to select two different immune cell types. In this example we will look at differences between two types of memory T cells: memory CD4 T-cells vs. memory CD8 T-cells.
31. Toggle between the three types of hierarchical visualizations: treemap, enclosure diagram, and Voronoi treemap. Finish by selecting the enclosure diagram tool.
32. The Hierarchy tool provides many of the same parameters: ‘Depth’, ‘Subgroup’, ‘Color map’. Two additional parameters are available for users to customize the plot:

a. Value: allows the user to display results in either the continuous (‘score’) or binary (‘activity’) form.
b. Label opacity: users can modify the label opacity using the sliding bar at the bottom left of the plot.
33. Set parameters:

a. Depth: 3 (task)
b. Subgroup: memory CD4 T-cell vs. memory CD8 T-cell
c. Value: activity
d. Color map: blue-orange
e. Label opacity: 0.75
34. Hover your mouse over a plot element to view a pop-out window with summary statistics of the element.
35. Display multiple summary statistics.

a. Select the task, system, and/or subsystem elements of interest. In this example we will look at: i) ‘GPI ANCHOR BIOSYNTHESIS’, ii) ‘Phosphatidyl-inositol to glucosaminyl-acylphosphatidylinositol’, and iii) ‘Malate to pyruvate conversion’. This can be done in 2 ways: 1) locate and select the subsystem bubbles directly from the enclosure diagram, or 2) search for elements using the drop-down menu above the plot. (Figure 5A)
b. Click on the 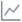 button to reveal the summary details of the selected tasks.
c. Use the ‘Subgroup’ parameter to toggle between summaries of individual groups (memory CD4 T-cell OR memory CD8 T-cell, Figure 5B-C) and differences between the two groups (memory CD4 T-cell vs. memory CD8 T-cell, Figure 5D).

**Figure 4:**
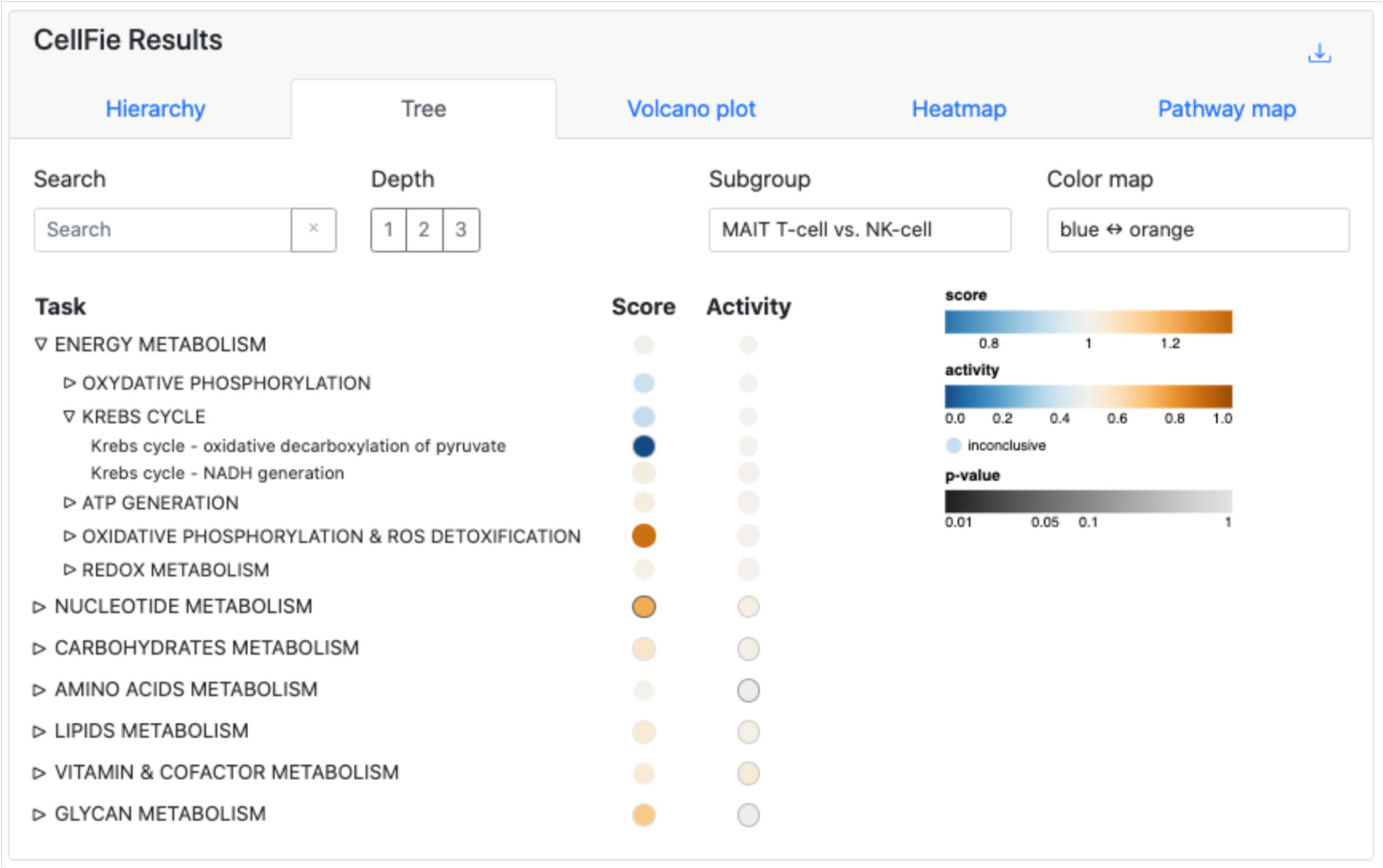
Treemap tool. The interactive treemap tool allows users to explore the CellFie Hierarchy. Users can click on various levels of the hierarchy to expand/shrink layers.

**Figure 5:**
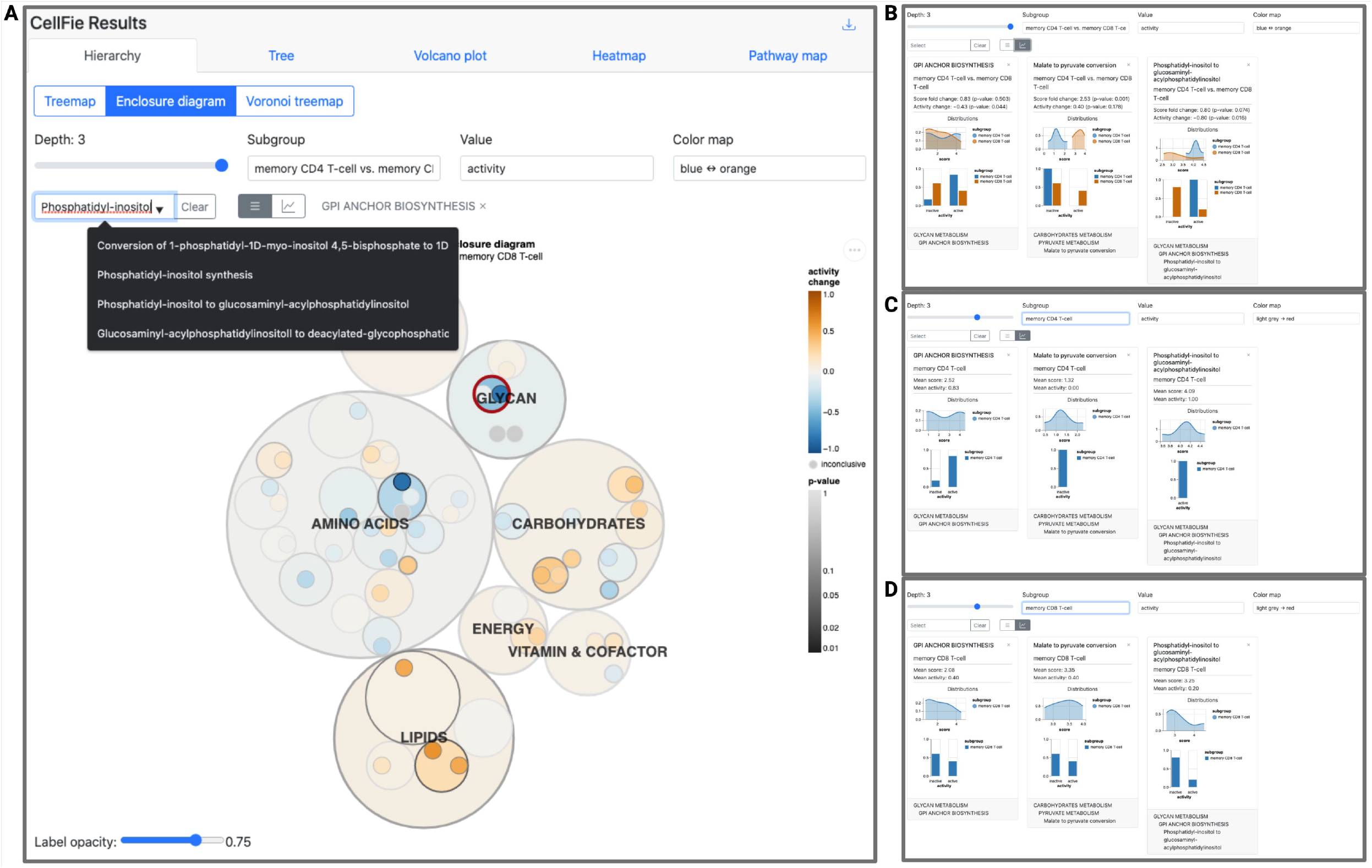
Hierarchy tool. Screenshots demonstrating use of the Hierarchy tool. A) Users can use the Search tool to select tasks, systems, or subsystems of interest. Here we have already selected the subsystem ‘GPI ANCHOR BIOSYNTHESIS’ (outlined in red in the plot and listed to the right of the search bar) and we are currently searching for the task ‘Phosphatidyl-inositol to glucosaminyl-acylphosphatidylinositol’. B-D) Summary statistics for 3 select CellFie elements showing the difference in activity scores (B) and the activity scores of the individual groups (C and D).

#### Optional

Users can save the hierarchy plot as or SVG or PNG using the 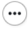 button to the right of the plot.

### Explore differences between cell types with the Volcano plot

**Timing:** minutes

ImmCellFie offers many interactive visualization tools for users to explore, including volcano plots. This tool allows users to visualize differences in CellFie scores (continuous) between phenotypic groups in the form of a volcano plot. Significantly upregulated and downregulated cell functions are colored in red and blue, respectively. Non-significant differences are shown in gray.

36. Click on the ‘Volcano plot’ tab from CellFie Results. **<u>Troubleshooting 4</u>**
37. Use the drop-down menus under ‘Select Subgroups to Compare’ to select two different immune cell types. In this example we will look at differences between 2 types of granulocytes: neutrophils vs. eosinophils.
38. Toggle between Depth 1, 2, and 3 of the CellFie hierarchy.

a. Observe the significant metabolic differences between the two granulocytes. In this example, we find significant differential activity of many metabolic tasks between neutrophils and eosinophils, especially tasks related to energy metabolism.
39. Explore the details of subsystems that show significant differential activity between the two cell types. (Figure 6)

a. While the volcano plot is on Depth 2 select the subsystems ‘OXIDATIVE PHOSPHORYLATION & ROS DETOXIFICATION’, ‘PYRIMIDINE CATABOLISM’, ‘AMINO SUGARS METABOLISM’, and ‘N-GLYCAN METABOLISM’. This can be done in 2 ways: 1) locate and select the subsystem bubbles directly from the volcano plot, or 2) search for the subsystems using the drop-down menu above the plot.
b. Click on the 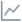 button to reveal the summary details of the selected subsystems.

**Figure 6:**
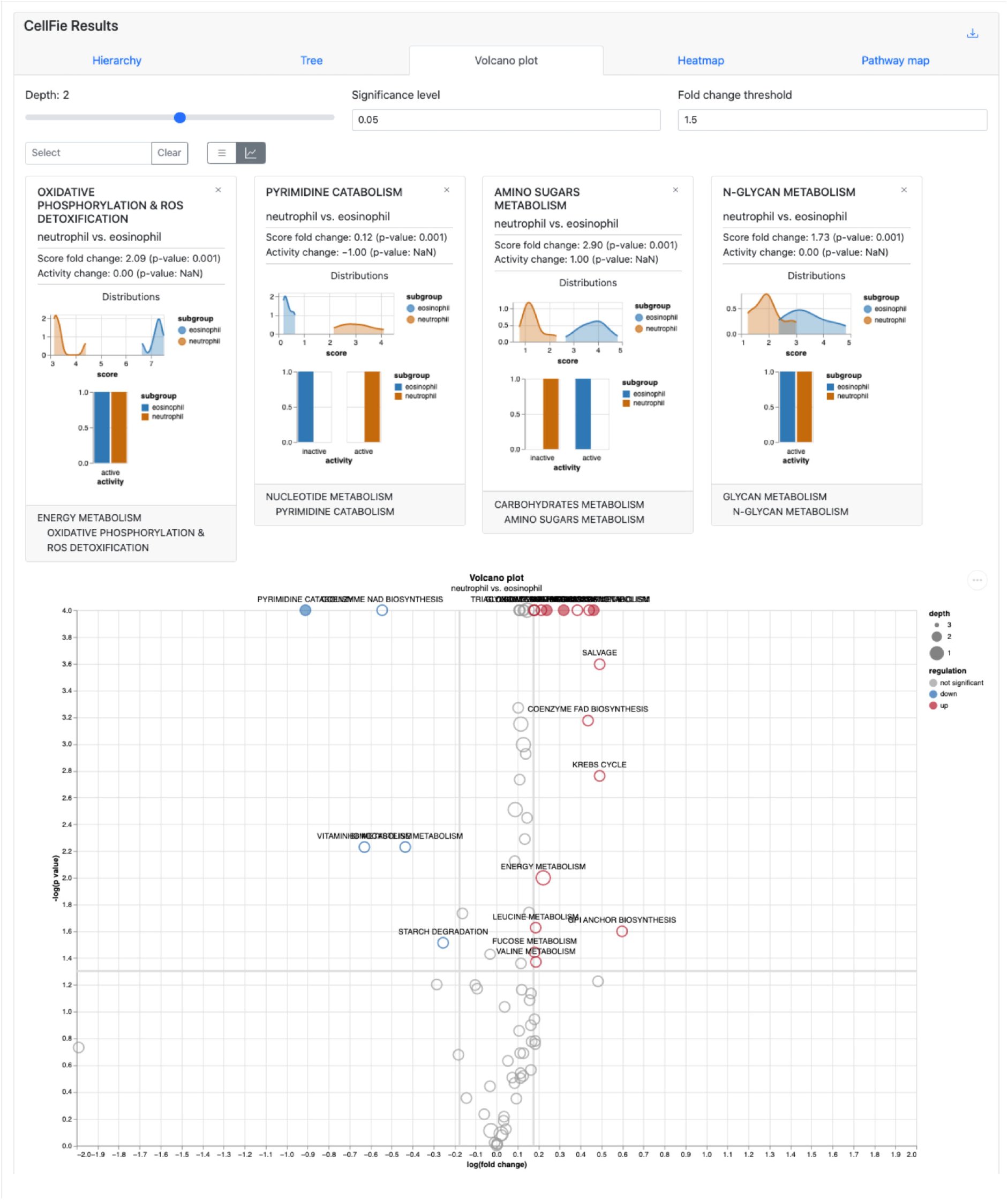
Volcano plot tool. Neutrophils and eosinophils are metabolically quite different. Here, we highlight 4 subsystems which show significant differential activity between the two cell types.

#### Optional

Users can change the significant level and fold change thresholds.

#### Optional

Users can save the volcano plot as SVG or PNG using the 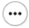 button to the right of the plot.

### Heatmap visualization of CellFie activity

**Timing:** minutes

ImmCellFie provides a Heatmap tool which generates a grouped heatmap of sample activity.

40. Click on the ‘Heatmap’ tab from CellFie Results.
41. Use the drop-down menus under ‘Select Subgroups to Compare’ to select two different immune cell types. In this example we will compare our custom groups Monocytes vs. Dendritic Cells.
42. In addition to the ‘Depth’, ‘Value’, and ‘Color map’ parameters you have seen in the previous tools, Heatmap allows users to sort samples by mean, median, or max using the ‘Sort by’ parameter.
43. Set ‘Depth’ to 2 (subsystem) and ‘Value’ to score (continuous). (Figure 7A)

a. Note that monocytes show increased activity in the top several subsystems.
44. Set ‘Depth’ to 3 (task), ‘Value’ to activity (binary), and ‘Color map’ to ‘yellow-green-blue’. (Figure 7B)

a. Note there are two tasks that are active in all dendritic cell samples and inactive in all monocytes: Synthesis of creatine from arginine & Conversion of lysine to L-2-Aminoadipate.

**Figure 7:**
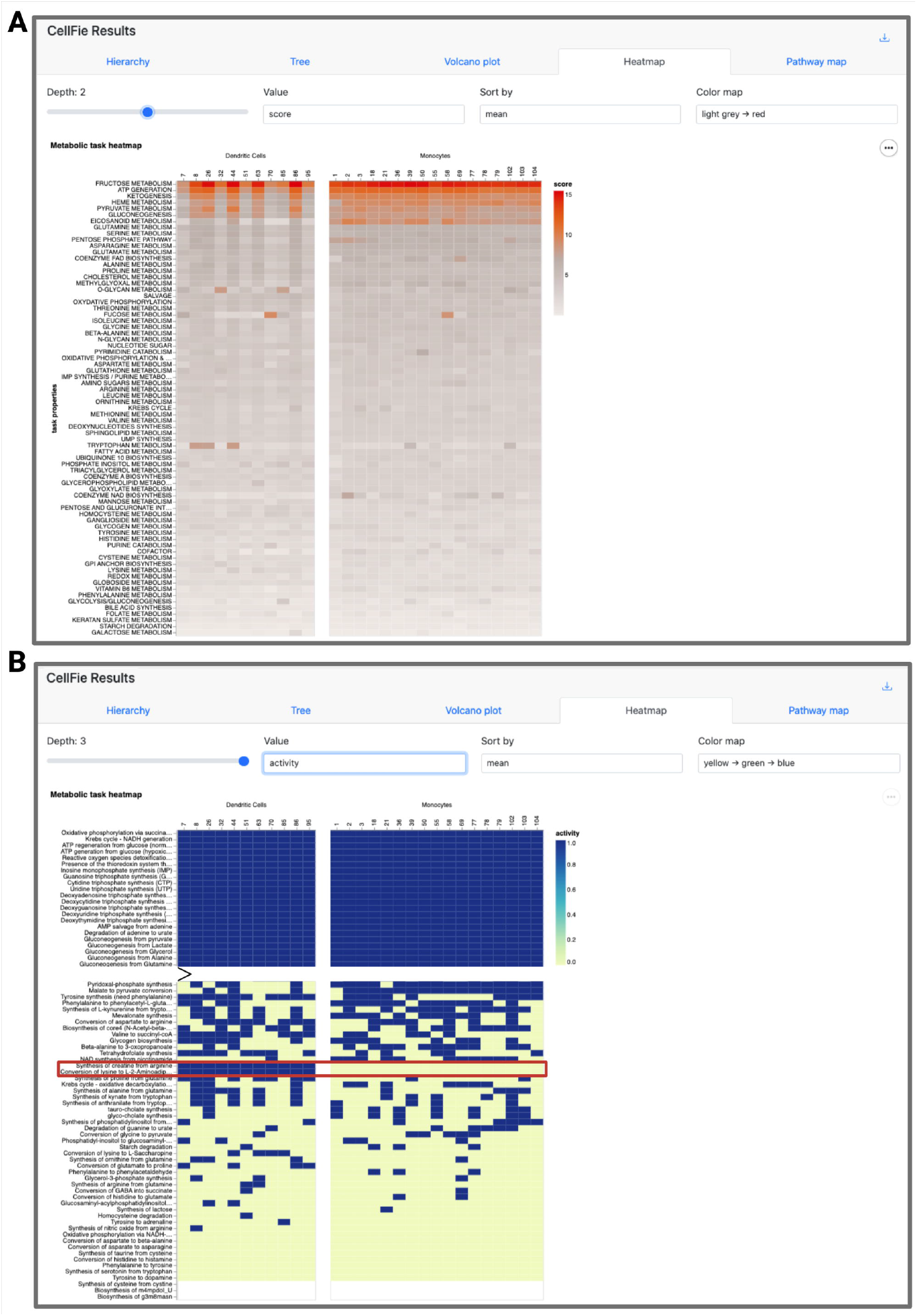
Heatmap tool. Heatmap showing metabolic differences between Dendritic Cells and Monocytes. A) Metabolic subsystem scores (continuous) of monocyte and dendritic cell samples. B) Metabolic task activity (binary) of the same set of samples.

#### Optional

Users can save the heatmap as SVG or PNG using the button to the 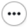 right of the plot.

### Visualizing metabolic activity of Escher pathway maps

**Timing:** minutes

This tool overlays reaction activity scores (column 6 of the detailScoring file) onto Escher (v.1.7.3) pathway maps (King *et al*., 2015) of CellFie’s 7 main metabolic systems: energy metabolism, nucleotide metabolism, carbohydrate metabolism, amino acid metabolism, lipids metabolism, vitamin and cofactor metabolism, and glycan metabolism. Users can visualize the pathway activity of single samples/groups or the differential pathway activity between two groups.

45. Click on the ‘Pathway map’ tab from CellFie Results.
46. Use the drop-down menu under ‘Select Subgroups to Compare’ to select two different immune cell types. In this example we will look at differences between naive B cells and memory B cells.
47. Use the drop-down menu under ‘Select map’ to toggle between the 7 metabolic system maps. Select the ‘Amino Acids Metabolism’ map.
48. The subgroup parameter can be used to view the pathway maps for individual groups (‘naive B cell’ OR ‘memory B cell’), or the differences between groups (‘naive B-cell vs. memory B-cell’). Set ‘Subgroup’ to ‘naive B-cell vs. memory B-cell’ to display the log2 FC between the mean reaction scores of the two groups. (Figure 8)
49. Use the drop-down menu under View to display the map in Full Screen. Alternatively, users can click on the 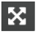 button to view the map full screen.
50. Users can customize the map using the Settings menu under View options.

a. Please refer to https://escher.readthedocs.io/en/latest/getting_started.html# sections 1.9 View options and 1.11 Settings for detailed information.

**Figure 8:**
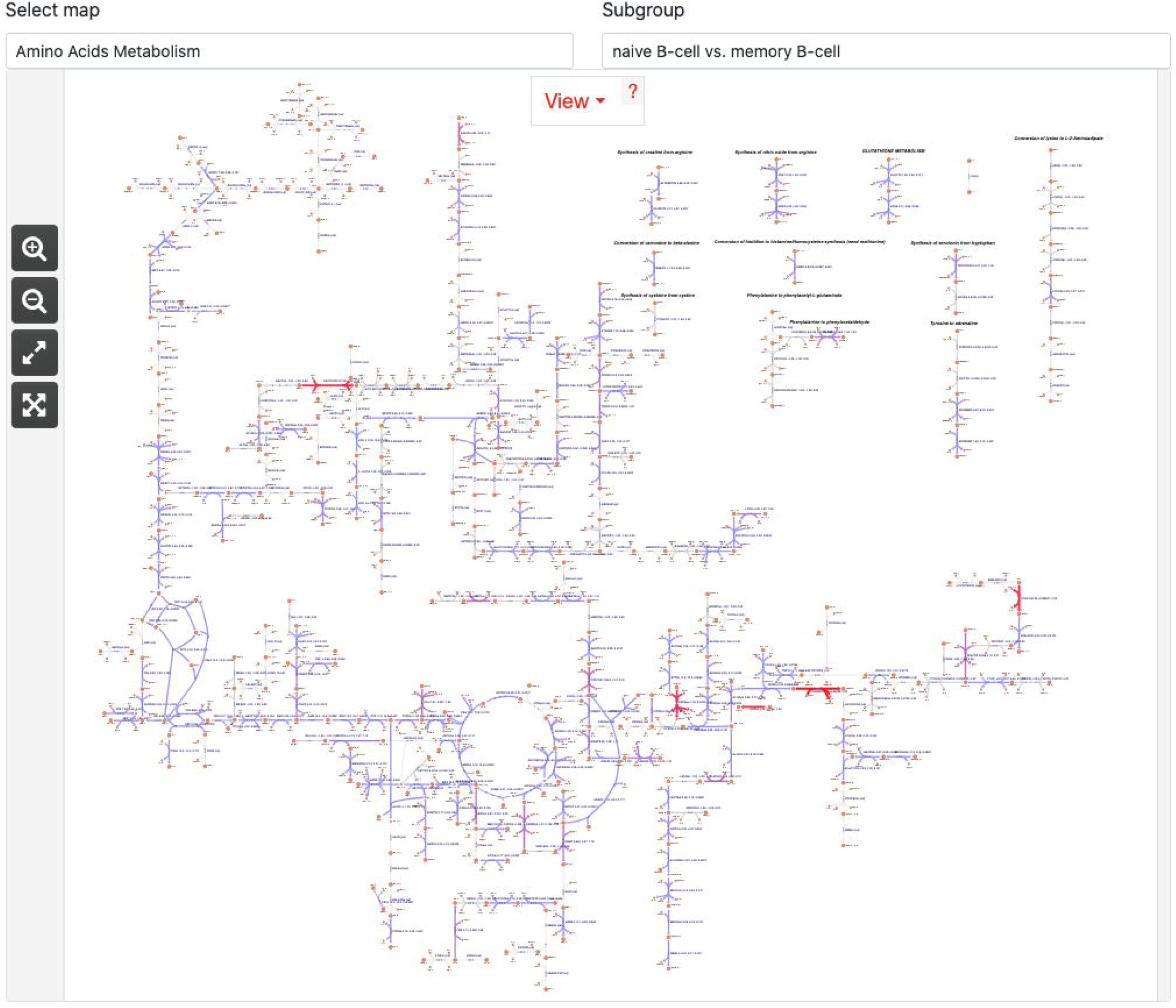
Pathway map tool. Escher pathway map of Amino Acid Metabolism differences between naive B cells and memory B cells. The Pathway map tool provides Escher pathway maps for the 7 systems of CellFie. The reaction activity scores of selected samples (using the subgroup parameter) are projected onto the reactions in the map and colored accordingly.

**Figure 9:**
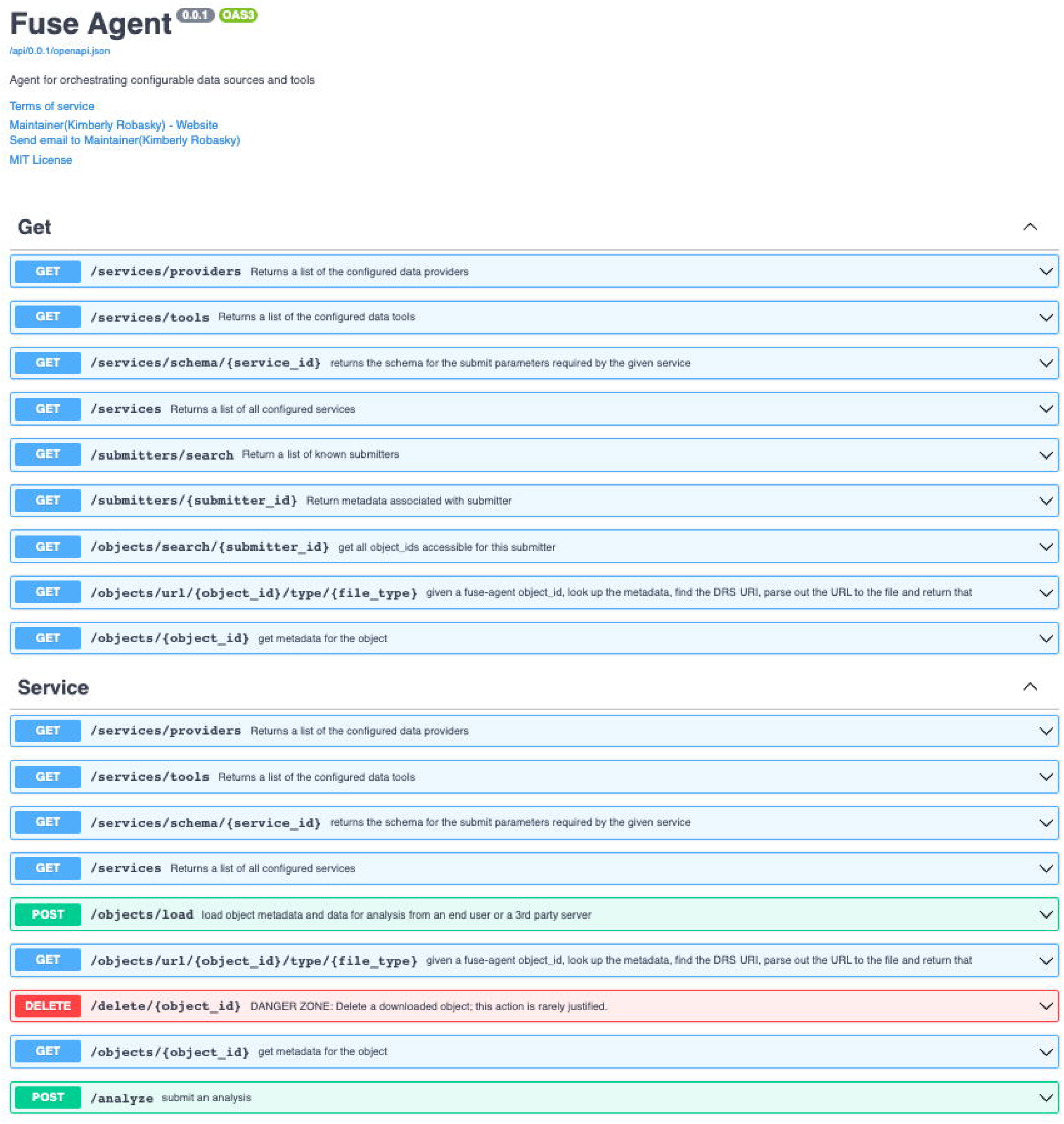
RESTful API. Screenshot of the RESTful, OpenAPI-compliant API system used to interface with ImmCellFie functions interactively over HTTP via notebooks (e.g., Jupyter, Google Colab).

## Expected outcomes

ImmCellFie is a user-friendly systems biology web tool for inferring metabolic activity from omics data and visualizing metabolic functionality with a variety of publication-ready plots. The CellFie algorithm can infer metabolic activity from transcriptomic or proteomic data for a variety of species including human, CHO, mouse, and rat. To facilitate exploration of immunological data, ImmCellFie supports direct data import from ImmuneSpace.org and the upload of local files. Once data has been uploaded to the ImmCellFie account, simple point-and-click features enable users to create custom metabolic analyses using CellFie’s thresholding parameters. The MTS output of CellFie provides a quantification of metabolic functions for each sample within the dataset. Due to the nature of how MTSs are calculated, scores should not be compared across tasks but only across samples. To facilitate intra-sample comparison of tasks, we provide the MTSs in a binary form which simply applies a task-specific threshold to determine whether the task is active or not. This form of binary scoring is referred to as activity. Users can use the ‘Downloads’ link at the top of the CellFie Results track to **batch-download** all the results and create custom downstream analyses using their program of choice.

ImmCellFie provides a suite of interactive analytical tools to visualize CellFie results. First, users can **create subgroups** within their dataset to facilitate conditional comparisons. This can be done manually by selecting which samples belong to designated groups, or semi-automatically based on information provided in the optional phenotypic file uploaded by the user. Here we used the immune cell category from the properties document to semi-automatically group samples based on cell type (Figure 3A). We also manually created custom subgroups of samples based on their lineage (Figure 3B). CellFie results can then be explored using ImmCellFie’s five interactive visualization tools: i) Hierarchy, ii) Tree, iii) Volcano plot, iv) Heatmap, and v) Pathway map.

CellFie tasks are organized into a hierarchical structure of metabolic pathways systems and subsystems, grouping tasks based on their biological functions. The Tree and Hierarchy tools enable users to explore and visualize scores at various depths within the CellFie hierarchy. The **Tree tool** allows users to interact with the architecture of the CellFie hierarchy linking tasks to their respective systems and subsystems, while including annotations of the score and binary activity results (Figure 4). Depth, subgroup, and color map parameters enable the user to toggle between levels of the hierarchy, individual subgroup scores and differential scores, and color schemes, respectively. While the Tree tool provides a means for users to explore the CellFie hierarchy to clearly understand the architectural links between tasks-systems-subsystems, it does not generate downloadable visualizations. The **Hierarchy tool** however provides publication-ready visualizations of the CellFie hierarchy in the form of a treemap, enclosure diagram, or Voronoi map. Here we set the parameters to explore the metabolic differences between two types of memory T cells: memory CD4 T cells vs. memory CD8 T cells. While the mature CD4+ and CD8+ subsets arise from a common progenitor, the CD4+/CD8+ double-positive (DP) cell which expresses both proteins simultaneously, they possess unique functions which are reflected in their unique metabolic signatures (Figure 5). Users can hover over elements of the plot to view summary statistics. Users can also select elements of the plot using their mouse or the search bar to select and display multiple summary statistics. Summary boxes include distribution plots of sample scores and binary activity. The summary of single subgroups will contain the mean score and activity of the group, otherwise the fold change and significant (p-value) metrics are provided. The bottom of the box also shows the parent term(s) of the selected CellFie function.

The **volcano plot tool** allows users to visualize differential metabolic scores between groups of samples. Here we compared two cell types from the granulocytes lineage: neutrophils vs eosinophils. Both neutrophils and eosinophils play an important role in the innate immune response, however each contain specific granules which impart unique metabolic demands on the cell. This can clearly be seen in the volcano plot of the two cell types (Figure 6). Sets of parameters allow the user to view differential activity at various levels within the CellFie hierarchy, as well as choose plot thresholds to define significant changes. Typically users would choose a significance level of 0.01 or 0.05, and a fold change threshold of 1.5 or higher. These functions enable immediate visualization of changes within the volcano plot, where significantly upregulated and downregulated cell functions are colored in red and blue, respectively. Similar to the Hierarchy tool, the interactive volcano plot allows users to select cell functions and display summary statistics of the subgroups.

ImmCellFie also provides a **Heatmap tool** which generates a grouped heatmap of subgroup sample scores. Here we used the tool to visualize the metabolic differences between our custom myeloid subgroups Monocytes and Dendritic Cells. Under disturbed homeostasis, monocytes are recruited to sites of inflammation and differentiate into distinct subsets of dendritic cells whose function is to stimulate effector T-lymphocytes (Wacleche *et al*., 2018). Users can customize the plot using the set of provided parameters. When the Depth parameter is set to 2 and we sort the subsystems by mean score, we can see that monocytes show increased activity in the top several subsystems, especially ‘EICOSANOID METABOLISM’ (Figure 7A). Interestingly, eicosanoids play an important role in the inflammatory response. We also observe distinct differences at the task level (Depth 1). In particular two tasks belonging to amino acids metabolism, synthesis of creatine from arginine and conversion of lysine to L-2-Aminoadipate, are active in all dendritic cell samples and inactive in all monocytes. Clearly, the cell types show distinct metabolic signatures reflecting their differences in lineage.

Lastly, ImmCellFie provides a **pathway map tool** to overlay cellular activity onto Escher maps of CellFie’s metabolic systems: energy metabolism, nucleotide metabolism, carbohydrate metabolism, amino acid metabolism, lipids metabolism, vitamin and cofactor metabolism, and glycan metabolism. Using the subgroup parameter, users can visualize single group activity or differential activity between two subgroups at the reaction level. Reactions are colored according to activity (blue-green-red). This enables users to quickly view changes related to specific reactions and enzymes within each metabolic system. Here we used the map to view metabolic differences between naive B cells and memory B cells. The Amino Acids Metabolism map shows clear differences in several reactions (Figure 8). To gain further insight into specific reactions and the enzymes of interest, users can hover over the reaction and navigate to the reaction page on the BiGG Models database.

In summary, the main features of ImmCellFie are: (1) a web-based platform enabling the inference of metabolic functions from omics data with the CellFie algorithm, (2) options to upload data from personal computer or import directly from ImmuneSpace.org, (3) an extensive suite of analytical and interactive visualization tools to explore results, (4) options to export raw CellFie results for further analyses or to generate of graphics for reports and publications using other software, (5) an intuitive user-friendly interface enabling scientists of any field to harness the power of GeMs, and (6) a RESTful, OpenAPI-compliant API for interfacing with all these functions interactively over HTTP via notebooks (e.g., Jupyter, Google Colab). The extendable design of the platform facilitates the addition of cell functionalities to expand the biological scope in future releases.

## Limitations

The CellFie tool is constrained by the availability of species-specific GeMs and generation of associated task modules. Therefore, the algorithm is currently compatible with a limited set of mammalian species and is limited to the inference of metabolic activity only. However, the modular design of the software can easily accommodate the integration of additional species and cellular functions as their GeMs and associated task modules become available.

The current state of CellFie does not consider pathway thermodynamics which are known to affect the pathway chemistries and fluxes in central metabolism(Flamholz *et al*., 2012; Noor *et al*., 2014). The incorporation of thermodynamics-based analysis with CellFie could be considered in a future release as an expanded feature.

The size of the dataset can pose issues for running the CellFie algorithm. This is especially relevant when attempting to run CellFie on single-cell datasets. To run large single cell datasets, users are advised to divide the dataset into smaller subsets and run them separately; however, this can impact the results since the local thresholding approaches are influenced by all samples included in the analysis. Thus, very large datasets with thousands of samples should be run locally.

ImmCellFie provides the user with the ability to easily generate multiple subgroups within the data to facilitate conditional comparisons. However, the suite of analytical and visualization tools only enables direct comparison between two groups and does not enable multi-group comparisons. In order to perform multidimensional analyses, the user should download the raw CellFie scores and use alternate programming tools.

## Troubleshooting

### Problem 1

Can I delete datasets?

### Potential solution

Users can request removal of datasets by navigating to the Data page and clicking on the “Remove” icon next to the dataset they wish to remove. This will open an email to the ImmCellFie development team requesting removal of the specified dataset.

### Problem 2

What should I do if the CellFie run is taking much longer than expected?

### Potential solution

This is most likely due to the size of the dataset (e.g. large single cell datasets). Upload the data in chunks and run them separately. Note that this will affect gene thresholding, however the effects should be minimal.

### Problem 3

Can I change the description of a dataset after I’ve already uploaded it and/or complete the CellFie run?

### Potential solution

Navigate to the Data Selection page, double-click on the description, and add/modify the description.

### Problem 4

Why did the CellFie run fail?

### Potential solution

There are 2 main reasons why CellFie runs might fail. First check the format of your input data and ensure the genes are in entrez ID form. Next check that the model/organism parameters correctly correspond to the organism genes used. We recommend googling several of your input genes to double-check they correspond to the correct organism.

### Problem 5

Why does the volcano plot x-scale not appear properly? (Figure 10A).

**Figure 10:**
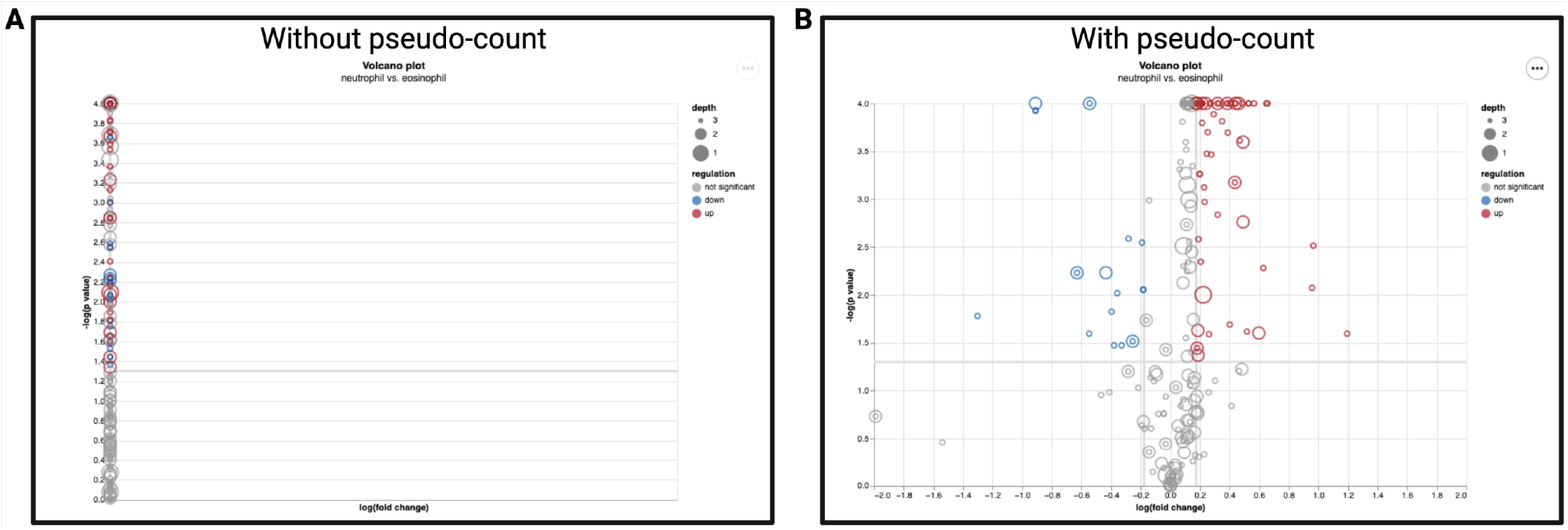
Troubleshooting the volcano plot. Example volcano plots showing the effect of adding a pseudo-count to the expression data.

### Potential solution

This issue arises due to how ImmCellFie handles the log form of zeros. To correct this issue, add a pseudo-count to the expression data and rerun CellFie (Figure 10B).

## Resource availability

### Lead contact

Further information and requests for resources and reagents should be directed to and will be fulfilled by the lead contact, Kimberly Robasky.

### Materials availability

The CellFie tool housed on ImmCellFie is freely available from: https://github.com/LewisLabUCSD/CellFie https://hub.docker.com/repository/docker/hmasson/cellfie-standalone-app Web application code: https://github.com/RENCI/fuse-dashboard-immcellfie (dashboard front-end code) https://github.com/RENCI/fuse-agent (dashboard back-end agent and API code)

Tools orchestrated by fuse-agent:

https://github.com/RENCI/fuse-tool-pca
https://github.com/RENCI/fuse-tool-cellfie
Datasets orchestrated by fuse-agent:

https://github.com/RENCI/fuse-provider-upload
https://github.com/RENCI/fuse-provider-immunespace

### Data and code availability

The example datasets and code generated during this study are included in this published article.

## Acknowledgments

This work was supported by generous funding from NIGMS (R35 GM119850 to N.E.L.) and NIAID (UH2 AI153029 to N.E.L. and K.R.) and the Novo Nordisk Foundation (NNF20SA0066621 to N.E.L.). The authors also thank Angela Wilden for preliminary analysis that inspired this work.

## Author contributions

K.R., H.O.M. and N.E.L. designed the study, conducted the analyses, and wrote the paper. A.R., D.B., J.R., A.T., L.B., S.S., M.W., Z.A.K., Z.L., B.P.K. developed the codes, tools, and visualization resources and contributed to analyses. L.C. coordinated efforts and facilitated code base issue tracking.

All authors have read and approved the work.

## Declaration of interests

The authors declare no conflicts of interest

---

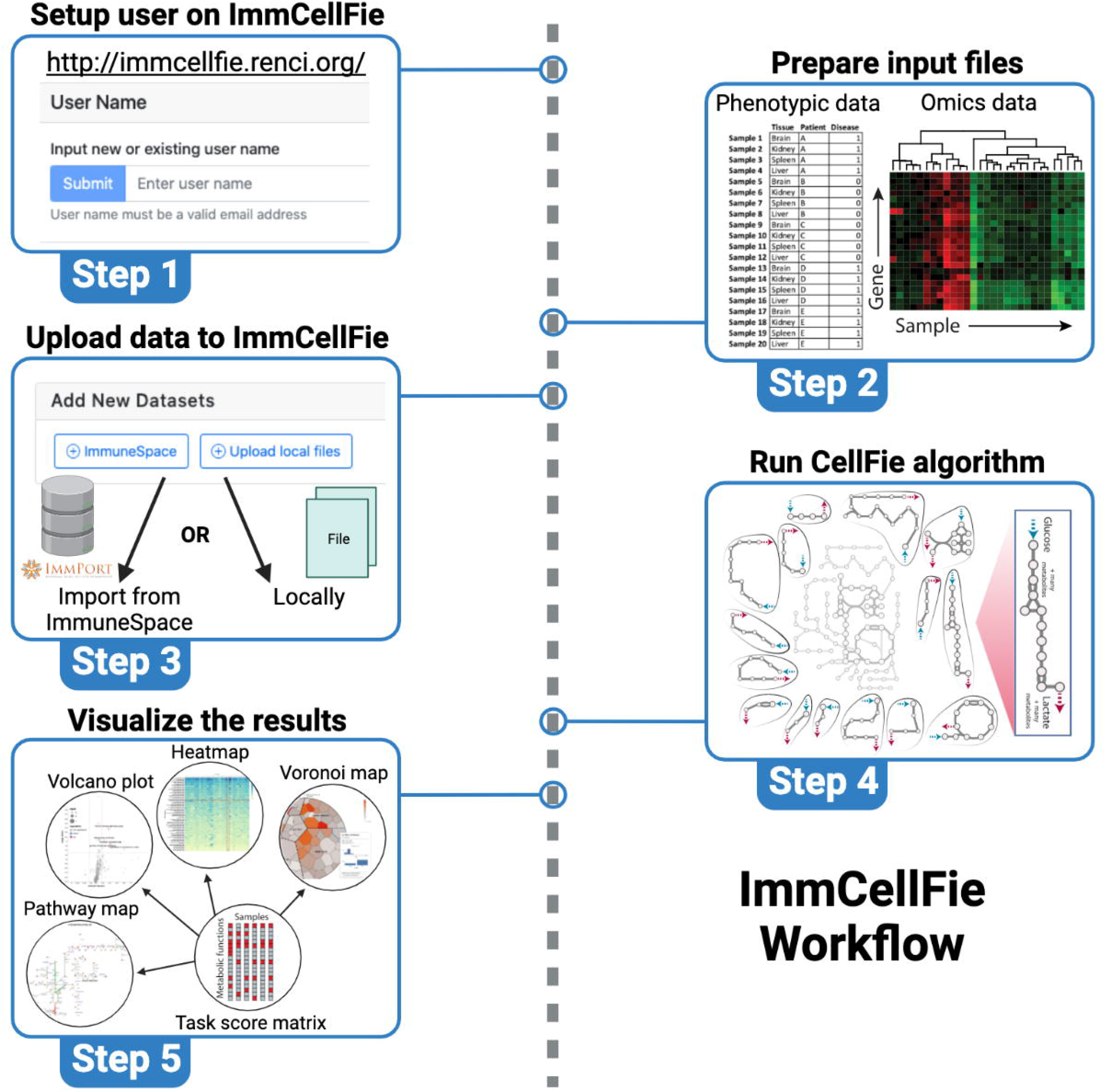

## Notes

### Competing Interest Statement

The authors have declared no competing interest.

